# Benchmarking Risk Predictions and Uncertainties in the NSCR Model of GCR Cancer Risks with Revised Low LET Risk Coefficients

**DOI:** 10.1101/2020.05.28.121681

**Authors:** Francis A. Cucinotta, Eliedonna Cacao, Myung-Hee Y. Kim, Premkumar B. Saganti

## Abstract

We report on the contributions of model factors that appear in fatal cancer risk projection models to the overall uncertainty in cancer risks predictions for exposures to galactic cosmic ray (GCR) in deep space. Annual GCR exposures to astronauts at solar minimum are considered. Uncertainties in low LET risk coefficients, dose and dose-rate modifiers, quality factors (QFs), space radiation organ doses, non-targeted effects (NTE) and increased tumor lethality at high LET compared to low LET radiation are considered. For the low LET reference radiation parameters we use a revised assessment of excess relative risk (ERR) and excess additive risk (EAR) for radiation induced cancers in the Life-Span Studies (LSS) of the Atomic bomb survivors that was recently reported, and also consider ERR estimates for males from the International Study of Nuclear Workers (INWORKS). For 45-y old females at mission age the risk of exposure induced death (REID) per year and 95% confidence intervals is predicted as 1.6% [0.71, 1.63] without QF uncertainties and 1.64% [0.69, 4.06] with QF uncertainties. However, fatal risk predictions increase to 5.83% [2.56, 9.7] with non-targeted effects. For males a comparison application to GCR using LSS or INWORKS lead to predictions of 1.24% [0.58, 3.14] and 2.45% [1.23, 5.9] without NTEs. The major conclusion of our report is that high LET risk prediction uncertainties due to QFs parameters, NTEs, and possible increase lethality at high LET are dominant contributions to GCR uncertainties and should be the focus of space radiation research.

## 1. Introduction

In this paper we benchmark the current uncertainties in estimating cancer risks from GCR exposures in the NASA Space Cancer Risk Model (NSCR). Because of the large uncertainties in high charge and energy (HZE) particle radiobiology and the small population of space workers, distinct methods are used at NASA to implement a radiation protection program compared to ground-based radiation workers. The basic approach is derived from recommendations by the National Council on Radiation Protection and Measurements (NCRP) [1-3], however we have developed an approach to make a rigorous uncertainty analysis [4-9], which has undergone external review by the National Research Council (NRC) [10] and NCRP [11].

The most recent analysis of GCR risks by the NSCR model [7-9] enjoys a significant reduction in overall uncertainty compared to our previous ones due to an improved treatment of the QF and DDREF and their possible correlations. Estimates of maximum relative biological effectiveness (RBE_max_) defined by the ratio of initial linear slopes determined at low dose and dose-rate for particles to γ-rays have been used in radiation protection to assign values of QFs. Values of RBE_max_ are highly dependent on the reference radiation used and their responses at low doses and dose-rates. The large values of RBE_max_ found in many experiments can be attributed in-part to the ineffectiveness of low doses or low dose-rates of γ-rays [8,9]. In addition, not all experiments have used either low dose-rates (<0.1 Gy/hr) or lower doses (<0.25 Gy) of γ-rays thus precluding RBE_max_ estimates. We have shown that assigning QF based on RBE’s for acute γ-ray exposures leads to a reduction in risk estimates and uncertainty.

The dominant uncertainties found in previous reports were the uncertainties in the parameters in the quality factor model and several uncertainties related to breakdown of the conventional risk assessment approach. Here the conventional approach using QFs only describe quantitative differences between heavy ions and other high LET radiation compared to a low LET reference radiation, while qualitative differences may occur. Furthermore because of the absence of epidemiology data for humans exposed to space radiation, the interpretation of data from experimental models are limited unless accurate extrapolation methods are developed. Previously we discussed two areas of possible qualitative differences, which are the higher lethality of high LET induced tumors compared to γ-rays or background occurring tumors, and the deviation from a linear response model due to non-targeted effects (NTE) [12,13]. We include estimates of their impact on GCR risk prediction in the updated analysis of this report.

The use of epidemiology data for populations exposed to γ-rays or other high energy photons has been the anchor to models that use RBE based QF’s to estimate space radiation risks. Here we note two recent studies provide updated analysis in the life-span study (LSS) of the survivors of the atomic-bomb explosions in Hiroshima and Nagasaki Japan in 1945 [14], and over 200,000 radiation workers from France, the United Kingdom and the United States [15-17]. The LSS analysis of Grant et al. [14] used revised dosimetry assessment and methods to correct for lifestyle factors compared to prior assessments [18-20]. An important finding by Grant et al. [14] is that for total solid cancer risk a linear-quadratic function provided an acceptable fit for males but not females. We consider these new data in our updated model denoted as NSCR-2020. For males, excess relative risk (ERR) based on linear coefficients are similar in these studies, however larger differences occur for tissue specific rates between the LSS and INWORKS studies. Therefore, we restrict our analysis to the grouped categories of all solid cancers and leukemia’s. We note that INWORKS uses mortality data, while we are using the LSS incidence data analysis converted to mortality predictions with current data for the US population. In addition, the INWORKS study does not provide data on the age or latency dependence of ERR or provide data on excess additive risk (EAR). We consider predictions for the US Average population, while updates for tissue specific predictions for never smokers will be reported in the future.

## 2. Model Development

We briefly summarize recent methods developed to predict the risk of exposure induced death (REID) for space missions and associated uncertainty distributions [7-9]. The instantaneous cancer incidence or mortality rates, λ_I_ and λ_M_, respectively, are modeled as functions of the tissue averaged absorbed dose *D*_*T*_, or dose-rate *D*_*Tr*_, gender, age at exposure *a*_*E*_, and attained age *a* or latency *L*, which is the time after exposure *L=a-a*_*E*_. The λ_I_ (or λ_M_) is a sum over rates for each tissue that contributes to cancer risk, λ_IT_ (or λ_MT_). These dependencies vary for each cancer type that could be increased by radiation exposure. However here we will group cancers into just two groups representing all total solid cancer risks and leukemia risk excluding chronic lymphocytic leukemias (CLL). The total risk of exposure induced cancer (REIC) is calculated by folding the instantaneous radiation cancer incidence-rate with the probability of surviving to time *t*, which is given by the survival function *S*_*0*_*(t)* for the background population times the probability for radiation cancer death at previous time, summing over one or more space mission exposures, and then integrating over the remainder of a lifetime [9]:

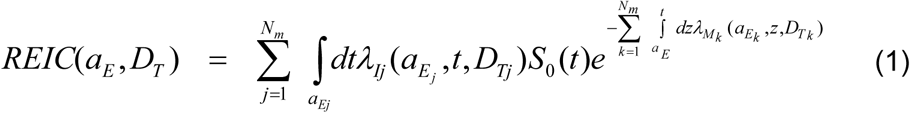

where z is the dummy integration variable. In equation (1), N_m_ is the number of missions (exposures), and for each exposure, j, there is a minimum latency of 5-years for solid cancers, and 2-years for leukemia assumed. Tissue specific REIC estimates are similar to equation (1) using the single term from λ_I_ of interest. The equation for REID estimates is similar to equation (1) with the incidence rate replaced by the mortality rate (defined below).

The tissue-specific cancer incidence rate for an organ absorbed dose, *D*_*T*_, is written as a weighted average of the multiplicative and additive transfer models, denoted as a mixture model after adjustment for low dose and dose-rates through introduction of the dose and dose-rate effectiveness factor (DDREF) and radiation quality through the R_QF_ factor related to the QF as described below:

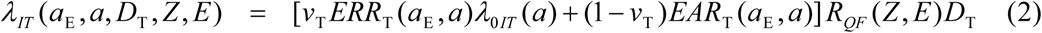

where *v*_*T*_ is the tissue-specific transfer model weight, *λ*_*0IT*_ is the tissue-specific cancer incidence rate in the reference population, and where *ERR*_*T*_ and *EAR*_*T*_ are the tissue specific excess relative risk and excess additive risk per Sievert, respectively. The tissue specific rates for cancer mortality *λ*_*MT*_ are modeled following the BEIR VII report [20] whereby the incidence rate of Eq. (2) is scaled by the age, sex, and tissue specific ratio of rates for mortality to incidence in the population under study:

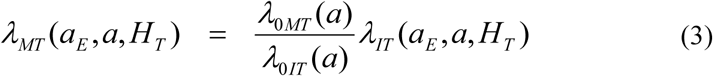

However, we also consider the possibility that high LET radiation increases incidence to mortality probabilities as described below. The U.S. lifetables from CDC [22] and cancer rates from SEER [23] with data collected from 2013-2017 are used to provide age and sex specific rates for survival of all causes of death and all solid cancer or leukemia excluding CLL. For cancer incidence we used the SEER delay-adjusted rates that accounts for delays that lead under-reporting of incidence data in the most recent year [23].

### 2.1 ERR and EAR Functions

Epidemiology studies for persons exposed to largely γ-radiation fit various statistical models to estimate ERR and EAR functions. The ERR and EAR functions used herein are for all solid cancers and leukemia risk excluding CLL. These functions depend on age at exposure, *a*_*E*_, and attained aged, *a*, using the parametric form:

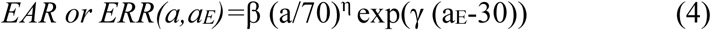

Values for the parameters in Eq. (4) from several reports [19,20] using similar functions are shown in **Table 1**, with calculations reported here using the recent Grant et al. results for solid cancer [14]. For leukemia risk we use the *ERR* and *EAR* functions from the BEIR VII report [21]. The BEIR VII used 60 instead of 70 in Eq. (4) which leads to a small difference in comparisons. The transfer model coefficient *v*_*T*_ have a large impact for individual cancers when background rates vary between the US and Japan, however predictions are less sensitive for total solid cancer risks. We use the mean value of *v*_T_=0.7 suggested by the BEIR VII report. For Monte-Carlo sampling we use a uniform distribution on (0,1) with *ERR* chosen if a random number if <0.7 and *EAR* chosen if not.

**Table 1.** Parameters of ERR and EAR functions described in the report from various sources [14,20,21].

| | $\beta_M$, Gy-1 | Attained Age power | Age at exposure | $\beta_M$, Gy-1 | Attained Age power | Age at exposure, y<sup>-1</sup> |
| --- | --- | --- | --- | --- | --- | --- |
|  | **ERR Model, Males** |  |  | **ERR Model, Females** |  |  |
| **RERF, 2007\*** | 0.35 [0.28, 0.43] | -1.65<br>[-2.1, -1.2] | -0.0186<br>[-0.0073, 0.0288] | 0.58 [0.4, 0.54] | -1.65<br>[-2.1, -1.2] | -0.0186<br>[-0.0029, 0.0073] |
| **BEIR VII** | 0.33 [0.24, 0.47] | -1.4 [-2.2, -0.7] | 0 | 0.57 [0.44, 0.74] | -1.4 [-2.2, -0.7] | 0 |
| **RERF, 2017** | - | - | - | 0.64 [0.52, 0.77] | -1.36<br>[-1.86, -0.84] | -0.0249<br>[-0.0139, -0.0357] |
| **RERF, 2017**<br>**LQ model** | 0.094 [<0.02, 0.23] | -2.7<br>[-3.58, -1.81] | -0.0249<br>[-0.0139, -0.0357] | - | - | - |
| **RERF, 2017\*\*** | *0.33 [0.25, 0.42]* | *-1.66*<br>*[-2.11, -1.2]* | *-0.0236*<br>*[-0.0128, -0.0343]* | 0.64 [0.52, 0.76] | -1.36<br>[-1.86, -0.84] | -0.0249<br>[-0.0139, -0.0357] |
|  | **EAR Models, Males per 10,000 PY per Gy** |  |  | **EAR Models, Females per 10,000 PY per Gy** |  |  |
| **RERF, 2007\*** | 43 [33,55] | 2.38 [1.9, 2.8] | -0.0274<br>[-0.0174, -0.0386] | 60 [51, 69] | 2.38 [1.9, 2.8] | -0.0274<br>[-0.0174, -0.0386] |
| **BEIR VII** | 22 [15,30] | 2.8 [2.15, 3.41] | 0 | 28 [22, 36] | 2.8 [2.15, 3.41] | 0 |
| **RERF, 2017** | - | - | - | 54.7 [44.7, 65.3] | 2.07 [1.64, 2.53] | -0.0357<br>[-0.0249, -0.0462] |
| **RERF, 2017**<br>**LQ model** | 21.7 [<-1.7, 47.7] | 2.89 [2.14, 3.68] | -0.0357<br>[-0.0249, -0.0462] | - | - | - |

\*90% Confidence intervals

The INWORKS study only provide a constant *ERR* estimate for males for cancer mortality caused by radiation [15-17]. We use their estimate for all solid cancers that assume a 5-year lag as in the LSS study with *ERR* = 0.37 per Gy with 90% confidence intervals [0.1, 0.67]. For all leukemia’s excluding CLL, *ERR*= 2.96 [1.17, 5.21].

### 2.2. Space Radiation Quality Factor

Our radiation quality approach uses concepts for particle track structure to devise a functional form that is fit to available radiobiology data to formulate a radiation quality factor. In this approach QF depends on particle charge number and kinetic energy or equivalent velocity. This is different from the International Commission on Radiological Protection (ICRP) approach where QFs are based on LET alone or similar the use of a radiation weighting factors that is dependent on particle type but not LET.

The hazard function in Eq. (1) is scaled to other radiation types and low dose-rates using a scaling factor denoted, *R*_*QF*_, which is made-up of a QF and DDREF. The *R*_*QF*_ is estimated from relative biological effectiveness factors (RBE’s) determined from low dose and dose-rate particle data relative to acute γ-ray exposures, which we denote as *RBE*_*γAcute*_ to distinguish from estimates from *RBE*_*max*_ based on less accurate initial slope estimates. The scaling factor is written [9]:

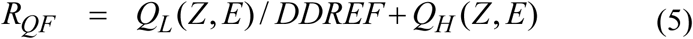

where

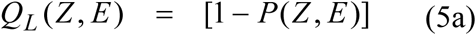

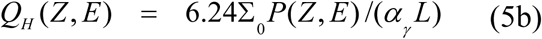

with the parametric function,

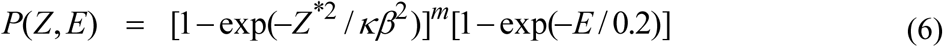

where *E* is the particles kinetic energy per nucleon, *L* is the LET, *Z* is the particles charge number, *Z\** the effective charge number, and *β* the particles speed relative the speed of light. The three model parameters (Σ_0_/α_γ_, κ and *m*) in Eq.’s (5-6) are fit to radiobiology data for tumors in mice or surrogate cancer endpoints as described previously [8,9,23,24]. Values and the cumulative distribution function (CDF) for Monte-Carlo sampling for the DDREF are described below. Distinct parameters are used for estimating solid cancer and leukemia risks based on estimates of smaller RBEs for acute myeloid leukemia and thymic lymphoma in mice compared to those found for solid cancers [9].

An ancillary condition is used to correlate the values of the parameter κ as a function of *m* as

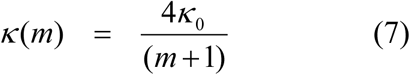

where κ_0_ is value for the most likely value *m*=3. In Monte-Carlo sampling by the model, conditional sampling is used where *m* is selected from a CDF followed by selection of *κ(m)*, which then distributed with a normal distribution with SD shown in **Table 2**.

**Table 2.** Parameters for central estimate of NASA quality factor (QF) parameters for solid cancer and leukemia risks [9].*

| Parameter | Solid Cancer | Leukemia |
| --- | --- | --- |
| **m** | 3±0.5 | 3±0.5 |
| **$\kappa$** | 624±69 (1000±250) | 624 ± 69 (1000±250) |
| **$\Sigma_0/\alpha_\gamma, \mu\text{m}^2 \text{ Gy}$** | (2897±357)/6.24 | (1750±250)/6.24 |
| **Non-Targeted Effects Parameters** |  |  |
| **$\eta_0/\alpha_\gamma, \text{Gy}^{-1}$** | 6 x10<sup>-5</sup> | - |
| **$\eta_1$** | 833± 200 (1000±150) | - |
| **$A_{\text{bys}}$** | 2000 $\mu\text{m}^2$ | |

\*Values in parenthesis are distinct values for light ions ( $Z \leq 4$ ).

A key assumption of the model is that the low ionization density part of a particle track is influenced by dose-rate effects as represented by the first term on the right-hand side of Eq. (5). However, the high ionization density part or so-called core of a particles track has no dependence on dose-rate as described by the second term on the right-hand side of Eq. (5). The low ionization density part of the track is high-energy δ-rays and they are expected to produce biological damage in a manner similar to low doses of γ-rays [23]. A dose-rate modifier is needed for the low ionization density track regions because model parameters are largely derived from radiobiological data at higher doses and dose-rates than those occurring in space, while a DDREF is used for this estimate.

The space radiation QF model corresponds to a pseudo-action cross section of the form which is of interest for fluence based risk prediction approaches,

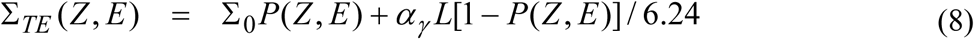

The Σ is denoted as a pseudo-biological action cross section for tumor induction in units of µm^2^ with the designation as “pseudo” given because time-dependent factors have been suppressed, which impact values for the cross-sectional area predicted by fits to the experiments.

The value of Σ_0_/α_γ_ estimated from mouse tumor studies [9] are shown in **Table 3**. We prefer to exclude mouse liver and Harderian gland values. The values for liver tumors are observed to be much larger in male compared to female mice, which needs to be further investigated. Harderian gland data are excluded here because this tissue does not appear in humans, and because related data are used as input for the other QF parameters so as not to put too much weight on a single model system. The CDF then follows a Gompertz equation with the parameters listed in **Table 3**.

**Table 3.** Cumulative distribution function (CDF) for model parameter (Σ_0_/α_γ_) determined by fits to data for heavy ions and fission neutrons*. Means and fits of CDF corresponding to values of **Table 3** using logistic or Gompertz equations are shown with best fit shown in bold font

| *Data sets* | *Mean, $\mu\text{m}^2 \text{ Gy}$* | *A* | *B, $\mu\text{m}^2 \text{ Gy}$* | *C* | *Adj R<sup>2</sup>* |
| --- | --- | --- | --- | --- | --- |
| *All solid cancer data* | (4728±1378)/6.24 | | | | |
| *Logistic Equation Fit* |  | **1.0±0.027** | **(2699±87)/6.24** | **-2.42±0.16** | **0.997** |
| *Gompertz Equation Fit* | | 1.0±0.02 | (2195±55.2)/6.24 | (1551±84.8)/6.24 | 0.987 |
| *Solid cancer excluding liver, and Harderian gland tumors* | (2897±357)/6.24 | | | | |
| *Logistic Equation Fit* | | 1.0±0.053 | (2483±110)/6.24 | -3.26±0.39 | 0.974 |
| *Gompertz Equation Fit* |  | **1.0±0.039** | **(2104±68.7)/6.24** | **(1109±110)/6.24** | **0.979** |

\*Parameters that result from fits of the logistic equation, $\text{CDF} = A/[1+((\Sigma_0/\alpha_\gamma)/B)^C]$ or Gompertz equation $\text{CDF} = A\exp[-\exp(-\Sigma_0/\alpha_\gamma - B)/C]$ to data for $(\Sigma_0/\alpha_\gamma)$ from mouse experiments for heavy ions and fission neutrons.

### 2.3. Dose and Dose-Rate Reduction Effectiveness Factor

Dose-rate is known to alter radiobiological effects at moderate to high doses. This is due to the effect of DNA repair, and modulation of tissue responses. For experimental studies with low particle fluences corresponding to less than one particle traversal per cell nucleus no dose-rate effect is expected at the molecular or cellular level, however tissue responses cannot be rule out. For spaceflight of duration of a few years or less dose protraction effects, which are distinct from dose-rate effects, are not considered. Estimating the DDREF for space exposures has the additional consideration compared to photon exposures on Earth because of the possible correlations between a dose-rate modifier and RBEs used to formulate a QF. Large RBE’s are often associated with large DDREFs.

In Cucinotta et al. [9], Bayesian analysis was used to model the PDF of uncertainty in the DDREF parameter for solid cancer risk estimates in a manner similar to that used in the BEIR VII report [20] were a prior distribution was estimated from the curvature in the Japanese Life-Space Study (LSS) and the likelihood function from radiobiology data. We denoted as Model A the prior distribution from the BEIR VII Report estimate for the LSS study using a log-normal distribution with a DDREF=1.3 and 95% confidence intervals (CI) of [0.8, 1.9]. However, Hoel [25] has argued that due to subjective assumptions made in the BEIR VII report a mean DDREF of 1.3 is found, while an analysis that considers a distinct dose range from the LSS data or one that includes downward curvature at higher doses due to cell sterilization effects finds a DDREF of 2 or more. Following Hoel’s analysis we used in Model B a mean DDREF of 2; however, uncertainties in this value were not modeled by Hoel. Here we assume a log-normal distribution with 90% confidence intervals of [1.2, 3] as a prior distribution for Model B based on the bounds described by Hoel [25]. Interestingly study of the curvature in dose response in the most recent LSS data by Grant et al [14] suggest a DDREF between 2 and 3 for males, while a DDREF∼1 for females. The effects of the large heterogeneous population in the study is a challenge for interpretation. In contrast experimental system offer a more precise method to estimate DDREF, however in less significant model systems and in some cases endpoints.

In our previous report we considered DDREFs from mouse solid tumor studies data where both γ-ray and high LET radiation were available. These data were used as the likelihood function for the Bayesian analysis as shown in **Figure 1** (upper panel). We did not consider ovarian and leukemia mouse data that was used by BEIR VII as appropriate for this analysis [8,9]. More recent experiments on heavy ion induction of colorectal and intestinal tumors (Suman et al., 2016) in mice did not provide other data to modify this aspect of the PDF of uncertainty for the DDREF because the γ-ray components of these experiments were limited, while dose responses for γ-rays in the recent Harderian gland experiments [26] were consistent with earlier data for Harderian gland tumors [27-29].

**Figure 1.**
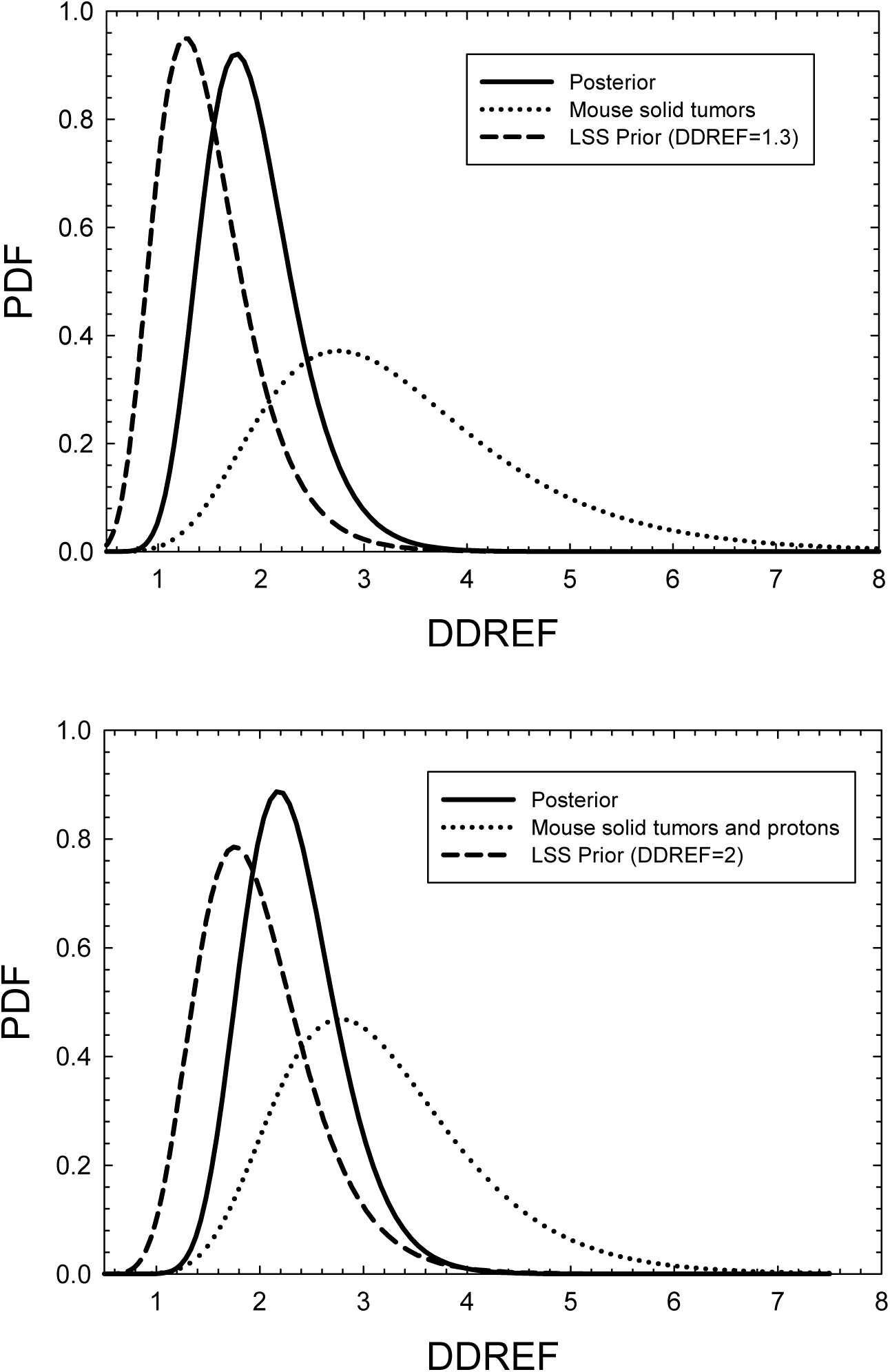
Bayesian analysis of probability distribution function (PDF) for the dose and dose-rate reduction effectiveness factor (DDREF). Upper panel uses prior distribution from the Japanese atomicbomb lifespan study (LSS) estimated in BEIR VII [20] with mean DDREF of 1.3 and likelihood function from mouse solid tumor studies with γ-rays. Lower panel uses mean DDREF of 2 as described in text for LSS study for the prior distribution, and likelihood function with mouse solid tumor studies with γ-rays and dose-rate studies for protons in surrogate cancer risk endpoints.

Values of DDREF’s estimated from high-energy proton experiments are of interest because the energy spectra of δ-rays more closely represent that of GCR compared to ^60^Co γ-rays [9]. We also surveyed published proton radiobiology data for tumors in animals and surrogate endpoints in cell culture models. Here we considered data comparing acute to low dose-rates, and analysis of curvature in acute dose response data to estimate a DDREF. In cell experiments several studies comparing high dose-rate to low dose-rates have been reported, which were summarized earlier [9]. DDREF estimates from proton experiments varied from 2.14 to 4.46 and strongly overlapped with estimates from solid cancers in mice exposed to acute and chronic doses of γ-rays. **Figure 1** (lower panel) shows the resulting PDF of the DDREF uncertainty in Model B which can be compared to our earlier publication for Model A [9].

### 2.4. Non-Targeted Effect Estimates

Non-targeted effects (NTE) have been shown to impact initiation, promotion and progression stages of tumorigenesis at low doses of high LET radiation [30-40]. Initiation processes impacted by NTEs include chromosomal exchanges, sister chromatid exchanges, gene mutation, and neoplastic transformation, which show a characteristic non-linear dose response at low particle fluences where less than one particle traverses a cell nucleus. A similar functional response provided an optimal global fit to the Harderian gland tumor study with several heavy ions [24]. Studies with La and Nb beams, which have action inactivation cross sections approaching or exceeding the cell nuclear area, suggest no cells directly traversed by these ions survive providing important evidence for NTE in this system. The use of chimera models where irradiated tissues absent of epithelial cells produce tumors after cell implant suggests changes to the micro-environment promote mammary tumors in mice [31]. Tissue are complex non-linear signaling systems contain multiple steady-states which can prevent excitability properties upon relaxation [41] leading to altered signaling and changes in proliferation and organization contributing to cancer development.

In our model we assume the TE contribution is also valid with a linear response to the lowest dose or fluence considered, while an additional NTE contribution occurs such a pseudo-action cross section is given by [24],

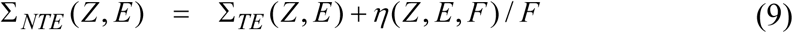

where *F* is the particle fluence (in units of µm^2^) and the η function represents the NTE contribution, which is parameterized as a function of *X*_*Tr*_*=Z*^*\*2*^*/β*^*2*^ as:

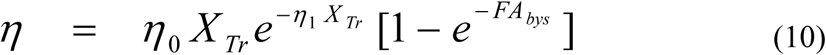

In Eq. (10) the area, *A*_*bys*_, reflects the number of bystander cells surrounding a cell traversed directly by a particle that receives an oncogenic signal. The RBE is related to the cross section by *RBE = 6.24 Σ/(LET α*_*γ*_) where α_γ_ is the γ-ray linear slope coefficient. Therefore, only the ratio of parameters η_0_/α_γ_ is needed for risk estimates.

The parameters η_0_/α_γ_ and η_1_ are estimated from low dose radiobiology experiments [24]. The second factor on the right-hand side of Eq. (9) describes the “turning on” of NTE at very low doses. The Harderian gland tumor model and chromosomal aberration experiments do not provide data of sufficiently low doses (<0.01 Gy) to determine at which dose or fluence level this occurs, and if it depends on radiation quality or the temporal patterns of irradiation. Therefore, the parameter *A*_*bys*_ is difficult to estimate. We note that its value is correlated with estimates of η_0_ at very low fluence since Eq. (10) here reduces to *η∼(η*_*0*_ *A*_*bys*_*) X*_*Tr*_ *exp(-η*_*1*_*X*_*Tr*_*)F*.

Several cell culture experiments were performed with α-particle irradiation which allow estimates of A_bys_. **Figure 2** shows an example for neoplastic transformation by 90 keV/µm α-particles with symbols with errors from experiments by Bettega et al. [40] and lines our model dose response function illustrating very low dose responses as *A*_*bys*_ varies. A possible turn-down of NTE at higher doses (>0.1 Gy) is ignored here because at these doses TE are expected to dominate. We find for a typical mammalian cell nucleus area of 100 μm^2^ that values of *A*_*bys*_ of 1000 to 2000 μm^2^ correspond to an NTE signal of about 1-cell layer and *A*_*bys*_ of 5000 μm^2^, a signal that propagates to about 2 cell layers from a directly hit cell. These areas suggest interaction distances of up to 50 microns from a directly traversed cell, and a reduction in NTE for doses below about 0.01 Gy (1 rad) where the NTE contribution decrease from a dose independent to linear response, while at higher doses (>0.1 Gy) TE dominate.

**Figure 2.**
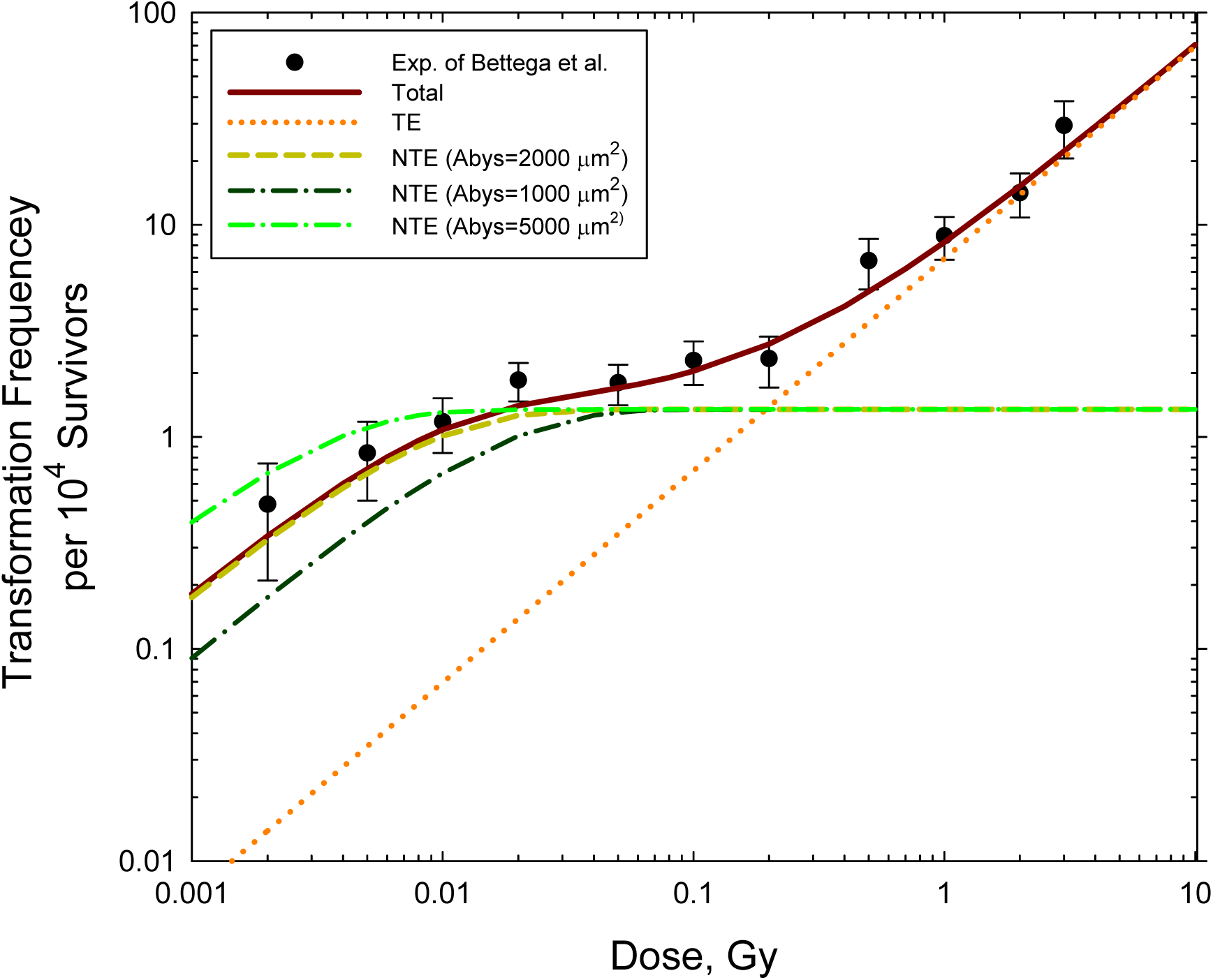
Dose response for neoplastic transformation of C3H10T1/2 cells by 90 keV/µm alpha particles. Experiments are from Bettega et al. [40]. Model shows characteristic mixed TE and NTE effects with NTE dominating at GCR type heavy ion doses (<0.1 Gy). Area of NTE estimated as A_bys_ ∼2000 µm^2^.

### 2.5 Implementation for GCR Exposures

GCR exposures include primary and secondary H, He and HZE particles, and secondary neutrons, mesons, electrons, and γ-rays over a wide energy range. We used the HZE particle transport computer code (HZETRN) with quantum fragmentation model nuclear interaction cross sections and Badhwar–O’Neill GCR environmental model to estimate particle energy spectra for particle type *j, φ*_*j*_*(Z,E)* as described previously [6, 42-46]. These methods agree with spaceflight data in low Earth orbit [4], in transit to Mars [44] and on the Mars surface [45] to within 15% for dose and dose equivalent. However larger differences between measurements and models occur for specific energy regions of particle spectra and therefore we have assigned a 25% variance for Z>2 and 35% for Z=1,2 ions.

For the TE model, a mixed-field pseudo-action cross section is formed by weighting the particle flux spectra, *φ*_*j*_*(E*) for particle species, *j*, contributing to GCR exposure evaluated with the HZETRN code with the pseudo-biological action cross section for mono-energetic particles and summing over all particles and kinetic energies:

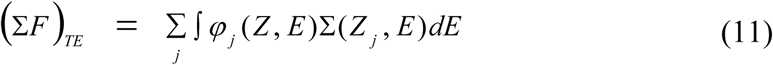

For estimates of NTEs to GCR exposures we assume: 1.) The probability that a bystander cell receives an oncogenic signal only occurs if the fluence is sufficiently high such that a nearby cell is traversed. 2.) The time dependence of the bystander signals is a few days or less such that interactions of bystander signals from different HZE particles can be ignored because of the low fluence in space. 3.) The probability that a bystander cell is transformed by a direct hit at a different time is small and can be ignored. Equations for the mixed-field pseudo-action cross section in the NTE model as folded with particle specific energy spectra as:

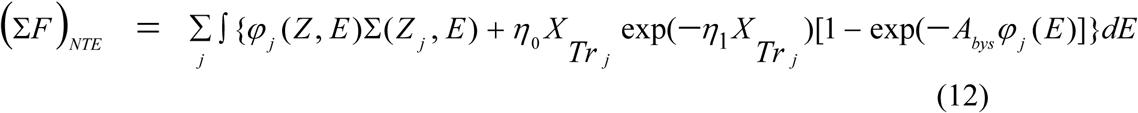

### 2.6 Sensitivity Study of Increased Tumor Lethality at High LET

We use the BEIR VII method to convert the LSS data from incidence to mortality predictions. This approach accounts to some extent to differences in conversion rate over time due to time-dependent differences in cure rates. However, these are still major questions on whether the quality of tumors produced depends on radiation quality. RBEs’ for both incidence and lethality in mice have not been reported, while differences in rates of metastasis and malignancies of tumors produce suggest difference do occur [47-54]. We estimated the effects of higher tumor lethality for HZE particles and neutrons in the following manner. An upper limit on the possibility of higher tumor lethality would be to use REIC estimates for REID estimates on space missions. However, this estimate would be too large due to the presence of low LET particles such as protons that make up a significant fraction of space radiation organ doses or the low ionization density part of HZE particle tracks which are a low LET radiation. To make a more realistic estimate of the effects of an increased lethality the cancer mortality rate is modified as [7]

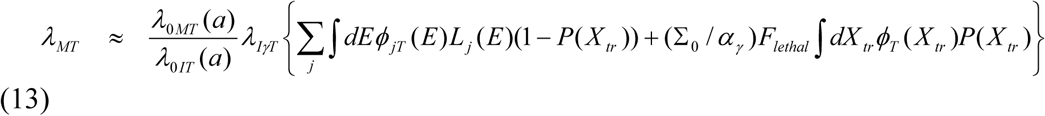

The first term in Eq. (13) dominates for low LET radiation and is not altered under the considerations of increased tumor lethality for highly ionizing radiation. The second term in Eq. (13) is increased by a tumor lethality fraction, *F*_*lethal*_ which estimates the increased lethality or rates of metastasis observed in mouse tumor induction studies with heavy ions. The second term in equation (13) has been reduced to be independent of the particle type, *j*, using the variable X_tr_=Z^*2^/β^2^ as described previously [6-9]. For the sensitivity study of *F*_*lethal*_, we considered a PDF to represent the uncertainty in the increased lethality for HZE particles and secondary charged particles from neutrons. The PDF is modeled as a normal distribution with several values considered assuming a 25% variance. We note that the RBE values for solid tumors considered previously were for tumor incidence and the sensitivity study of Eq.(13) is not used for leukemia risk estimates because there is no evidence for increased high LET mortality compared to low LET from mouse studies.

For the application of the NSCR model to space mission predictions, the energy spectra for each particle type, *j* of LET, *L*_*j*_*(E)* for each tissue, T contributing to cancer risk denoted as *ϕ*_*j*T_*(E)* is estimated from radiation transport codes. The particle energy spectra are folded with *R*_*QF*_ to estimate tissue specific REIC or total REID values. For calculations for a fluence *ϕ*_T_*(Z,E)* and absorbed dose, *D*_T_*(Z,E)* of a particle type described by *Z* and *E*, the Hazard rate is

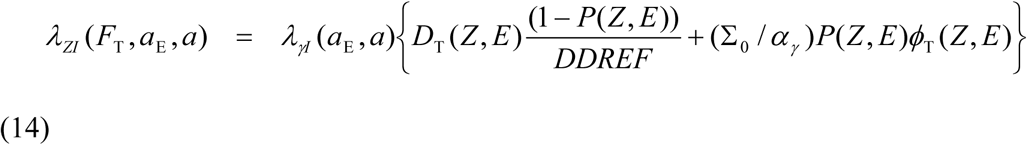

where λ_γI_ is the inner bracketed terms that contains the *ERR* and *EAR* functions for individual tissues. As described previously [9] calculations are made using models of the GCR environments and radiation transport in spacecraft materials and tissue, which estimate the particle energy spectra, *ϕ*_*j*_*(E)* for 190 isotopes of the elements from *Z*=1 to 28, neutrons, and dose contributions from pions, electrons and γ-rays.

The calculation is simplified by introducing the fluence spectra, *F(X*_*tr*_) where X_tr_= Z^*2^/β^2^, which can be found by transforming the energy spectra, *ϕ*_*j*_*(E)* for each particle, *j* of mass number and charge number, *A*_*j*_ and *Z*_*j*_ respectively as:

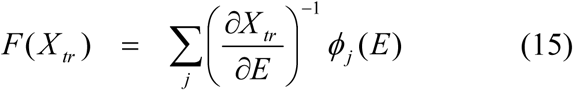

where we evaluate the Jacobian in equation (12) using the Barkas form for the effective charge number given by

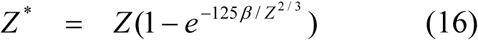

This transformation allows the REID calculation to occur with the tissue specific Z*^2^/β^2^ spectra for light and heavy ions rather than the individual Z and E spectra.

### 2.7. Summary of Parameter Uncertainty PDFs

For the various parameters that enter into the model PDFs that are estimated from experimental data and model comparisons to represent plausible ranges of values. The uncertainty in the ERR and EAR parameters are taken directly from the publications noted above. Values showed modest skewing and therefore we used a normal distribution for each parameter with standard deviations (SD) estimated from the publications. Our recent report [9] used solid tumor data in mice directly to model the value and PDF for the parameter Σ_0_/α_γ_, where the PDF is represented by the Gompertz equation (**Table 3**). Bayesian analysis is used to model the uncertainty in the DDREF parameter. The BEIR VII Report estimate [17] for the Japanese Life-Span Study (LSS) study of DDREF=1.3 with 95% confidence intervals (CI) of [0.8, 1.9] was used as the prior distribution, which is updated using Bayes’ theorem with the likelihood function represented by a log-normal distribution. The resulting posterior distribution has a mean value of 1.88 with 95% CI of [1.18, 3.0]. For the central values of REID estimates for space missions discussed below we continue to use the value DDREF=2, however the posterior distribution is used to represent the PDF for the DDREF uncertainty in the analysis described here, which is also fit to a log-normal distribution (**Table 4**). Other parameters are similar to early versions of NSCR.

**Table 4.** Summary of Probability Distribution Function (PDF) used for various terms and their parameters.

| Model Term | CDF for Monte-Carlo Sampling | Parameters |
| --- | --- | --- |
| ERR, EAR functions | Normal | SD for parameters from source with additional 10% variance added. |
| DDREF | Log-Normal | GM=0.83; GSD=1.22 |
| QF parameter, $\Sigma_0$ | Gompertz Equation | See Table 3. |
| QF parameter, $m$ | Normal | SD=0.333 |
| QF parameter, $\kappa$ | Conditional sampling on $m$, followed by Normal | SD=0.25 |
| NTE parameter, $\eta_0 \times A_{bys}$ | Normal (See Table 2). | SD=0.333 |
| NTE parameter, $\eta_1$ | Normal (See Table 2) | SD=0.333 |
| Fluence $Z=1,2$ | Normal | $M=1$; SD = 0.35 |
| Fluence $Z>2$ | Normal | $M=1$; SD = 0.25 |

**Table 4.** % REID and uncertainties for 45 y old female and male astronauts for annual GCR exposure near solar minimum with 20 g/cm^2^ aluminum shielding.

| Uncertainties included | %REID | 95% Confidence Intervals |
| --- | --- | --- |
| **Females** |  |  |
| **LSS and Transfer** | 1.53 | [1.38, 1.63] |
| **LSS, Transfer and DDREF** | 1.19 | [0.98, 1.43] |
| **LSS, Transfer, DDREF, and QF** | 1.6 | [0.71, 4.06] |
| **LSS, Transfer, DDREF, QF and space organ dose** | 1.64 | [0.69, 4.35] |
| **Males** |  |  |
| **LSS and Transfer** | 1.14 | [1.05, 1.21] |
| **LSS, Transfer and DDREF** | 0.92 | [0.77, 1.09] |
| **LSS, Transfer, DDREF, and QF** | 1.21 | [0.59, 2.91] |
| **LSS, Transfer, DDREF, QF, and space organ dose** | 1.24 | [0.58, 3.14] |

## 3. Results and Discussion

For all calculations we considered 45-y US Average male or female astronauts, and assumed an average spacecraft shielding amount of 20 g/cm^2^ of aluminum for annual GCR exposures near solar minimum. Predictions of REID for other shielding materials and amounts, and for other ages at exposure were considered previously for the NSCR-2012 model [6]. In **Table 4** and **Figure 3** we show %REID predictions and 95% confidence intervals (CI) for various uncertainty inclusions. We use the value for Σ_0_/α_γ_ that excludes liver and Harderian gland data. Use of these values would increase estimates by ∼30%. For males we use the linear fits which is consistent with the main result for females in the LSS report. However, Grant et al. [14] found an improved fit for males using a linear-quadratic dose response model, which is discussed below. The inclusion of the DDREF uncertainty tends to lower REID predictions, however it is not a large effect since the QF has been defined such that the track core term is independent of dose-rate.

**Figure 3.**
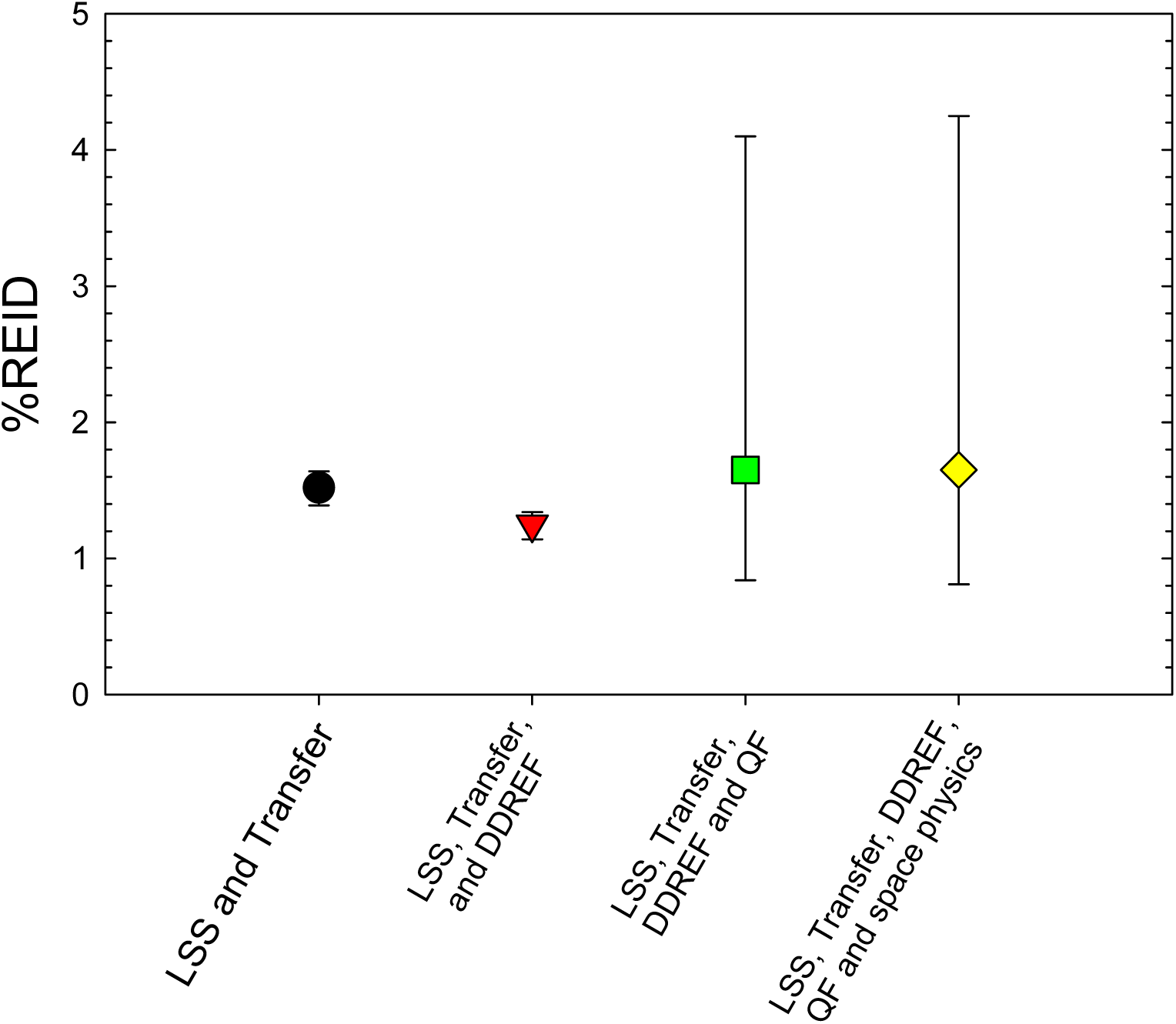
%REID predictions for 45-year old Females with different model uncertainties considered.

The uncertainties in the quality factor parameters dominate the uncertainty and shift average %REID predictions to higher values. The ratio of the upper 95% CI to the average value is <2.8. In **Table 5** we show a breakdown of the QF uncertainty for female astronaut %REID predictions. Result show that the Σ_0_/α_γ_ uncertainty makes the largest contribution followed by the uncertainty in the κ parameter. **The value of *m* which is highly constrained based on previous analysis [9,13] plays only a minor role, which suggests the QF has been reduced in effect to a two-parameter model**.

**Table 5.** Comparison of 45-y old males %REID predictions using LSS and INWORKS coefficients in multiplicative risk model or mixture model (LSS linear-quadratic).

| Low LET Model | % REID | 95% CI |
| --- | --- | --- |
| **LSS Linear (DDREF=2)** | 1.24 | [0.58, 3.14] |
| **LSS linear-quadratic fit using linear term only (DDREF=1)** | 0.6 | [0.4, 1.18] |
| **INWORKS (DDREF=1)** | 2.45 | [1.23, 5.9] |

For males we made a comparison of the LSS models to the INWORKS models (**Table 6**). Here the INWORKS analysis considers only ERR and does not consider any age or time after exposure parameterizations for the adult worker populations in the study. The life-table representing age dependent background cancers and deaths due to competing risks thus represent the only time dependent factors in this INWORKS application, while the LSS studies provide time dependencies as described by Eq. (4). It would be difficult to assess the higher prediction of the INWORKS rates to a single factor. Other differences include chronic versus acute exposure, contributions to organ doses from neutrons, and the effects of the various background populations in the different studies. In addition, we are using the incidence data from LSS converted to mortality using Eq. (3), while mortality data is used directly in the INWORKS study. Incidence to mortality conversion varies with time period, host country, and individual subjects health care all of which can impact the result.

**Table 6.** Sensitivity of %REID predictions on uncertainties in parameters of the cancer risk cross section for 45-y old females for annual GCR exposure near solar minimum. All non-QF uncertainties for a conventional model included.

| Uncertainty after Model Parameter Eliminated | %REID | 95% CI |
| --- | --- | --- |
| **m** | 1.6 | [0.7, 4.2] |
| **κ** | 1.42 | [0.77, 3.2] |
| **Σ<sub>0</sub>/α<sub>γ</sub>, μm<sup>2</sup> Gy** | 1.46 | [0.73, 2.84] |
| **All uncertainties** | 1.64 | [0.69, 4.35] |

### 3.1. Uncertainties due to Qualitative Differences

The application of radiation quality factors accounts for quantitative differences between radiation types, however does not represent possible qualitative differences in cancer risk for different types of radiation. Possible qualitative differences suggested by past studies include non-targeted effects, and differences in tumor lethality not estimated with RBEs based on tumor incidence or surrogate markers. Differences in latency and genetic background on radiation quality are also possible however there is insufficient data to make numerical estimates in this area.

Several reports [47-53] have suggested that HZE particles and neutrons could produce more lethal tumors compared to tumors produced by low LET radiation or background tumors. For low LET radiation there is an implicit assumption made by epidemiology models that the tumors induced by radiation are similar to background tumors in a population. This assumption is consistent with the multiplicative risk model, and also based on lack of information to make an alternative assumption. Using the sensitivity analysis method described earlier [7-9] suggests that increases in tumor lethality for HZE particle and neutrons compared to background or low LET tumors as suggested by animal studies could substantially increase REID and uncertainty estimates.

**Table 7** shows predictions for increased lethality of tumors at high LET using the formalism described above. In **Table 8** we show predictions with NTE. Both important high LET effects shift REID predictions dramatically to higher values. In **Figure 4** we compare probability distributions for the different models. NTE suggest a much higher level of concern than increased lethality. Also there is a much larger body of evidence that NTE’s will contribute to the mutation and instability at low doses of high LET, while few studies have directly investigated tumor quality effects.

## 4. Conclusions

Past NAS [54] and NCRP [1-3] reports where highly influential in stressing the importance of understanding the radiobiology of heavy ion and other high LET radiation, while not blindly assuming GCR risks are easily projected form low LET observations of risk. In-fact NCRP Reports No 98 and No 132 were intended only for low Earth orbit. NCRP Report 132 relied on the uncertainty assessment in NCRP Report 126 [2,3]. This report used largely subjective methods to estimate uncertainties in low LET radiation epidemiology including uncertainties in data collection, bias, errors in organ dose assessments of the atomic bomb exposures, future projections of immature data sets and statistical uncertainties. Larger uncertainties were estimated for estimating dose-rate effects for low dose and dose-rate exposures and transfer models that chose between EAR versus ERR. Since that time low LET radiation epidemiology data has matured to a great extent, while only modest uncertainties are estimated for total solid cancer and leukemia risks.

**Table 7.** Effect of increased tumor lethality for high LET radiation on %REID predictions for 45-y old females for annual GCR near solar minimum.

| Increased High LET Lethality coefficient | % REID | 95% CI |
| --- | --- | --- |
| 0 | 1.64 | [0.69, 4.35] |
| 20% | 1.86 | [0.71, 5.11] |
| 40% | 2.06 | [0.74, 5.89] |
| 60% | 2.24 | [0.77, 6.62] |

**Table 8.** Predictions of 45-y old females %REID predictions for average GCR conditions with addition of non-targeted effects.

| $A_{bys}, \mu m^2$ | % REID | 95% CI |
| --- | --- | --- |
| 0 | 1.64 | [0.69, 4.35] |
| 1000 | 3.67 | [1.68, 6.82] |
| 2000 | 5.83 | [2.56, 9.7] |
| 5000 | 12.2 | [5.1, 19.0] |

**Figure 4.**
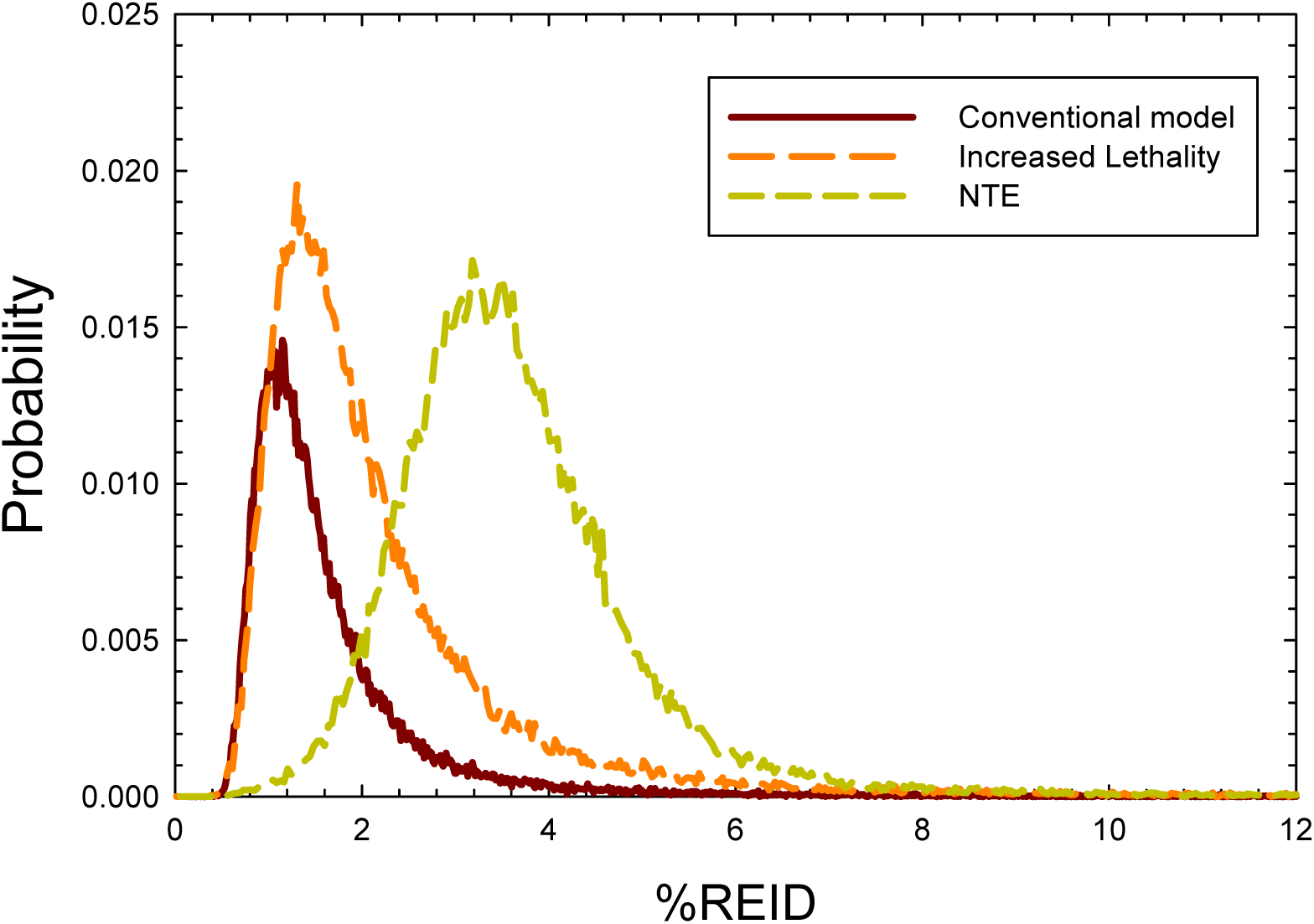
Probability distribution functions for 45-y old females in the conventional model and two predictions of the impact of increased lethality and NTEs.

The concordance in excess relative risks per Gy found between the LSS study and INWORKS suggest an agreement of about a factor of 2, however there are many differences in the makeup and maturity of the studies. The lack of an age and latency parameterization in INWORKS is a limitation in comparisons that use applications of these results. For tissue specific cancer risks larger differences occur. For example, Preston et al. [20] find many specific tissues have significant increases attributed to radiation exposure, while Richardson et al. [17] find non-significant results for many of the same tissues including colon, brain, liver, and bladder which makeup important contribution in the LSS study. The reason unknown but could be due to the lower doses in the INWORKS cohorts or differences in genetic or host environmental factors. In this report we considered only total solid cancer and leukemias excluding CLL. The treatment of tissue specific risks and update on background rates for never-smokers will be considered in a future report.

High LET related uncertainties in QF parameters, NTEs and tumor lethality were shown to dominate uncertainties. Track segment irradiation studies with heavy ions are need to reduce uncertainties in QF parameters. The dichotomy in the κ for light and heavy ions is likely due to the higher effectiveness of lower energy δ-rays (<5 keV), which has a larger impact of light ions. This effect will be addressed in future version of NSCR. Several recent reviews have noted the importance of NTE’s for high LET radiation and the supra-linear dose responses produced by NTE’s at low dose can substantially increase RBE estimates and skew PDFs for cancer risk estimates. Similar reports [32-42] have suggested that HZE particles and neutrons could produce more lethal tumors compared to tumors produced by low LET radiation or background tumors, which is a qualitative difference not accounted for in current risk estimates. For low LET radiation there is an implicit assumption made by epidemiology models that the tumors induced by radiation are similar to background tumors in a population. This assumption is consistent with the multiplicative risk model, and also based on lack of information to make an alternative assumption. The potential role for NTEs is the largest uncertainty found in this study. NTEs are supported by many mechanistic studies, however are sparse for dose response modeling. Studies are needed over the low dose range (0.001 to 0.05 Gy) in mouse or other small animals. Also, the use of high Z ions such as Nb, La, Au or Pb with ranges of a few cm or more in tissue are recommended because here directly traversed target cells have a high probability of cell kill. Therefore, tumors observed would likely directly NTEs.

## References

1. NCRP. Guidance on radiation received in space activities. National Council on Radiation Protection and Measurements. NCRP Rep. 98: Bethesda, MD; 1989.

2. NCRP. Recommendations of dose limits for low earth orbit. National Council on Radiation Protection and Measurements. NCRP Report 132: Bethesda MD; 2000.

3. NCRP, Uncertainties in fatal cancer risk estimates used in radiation protection. National Council on Radiation Protection and Measurements Report 126: Bethesda MD; 1997.

4. Cucinotta FA, Schimmerling W, Wilson JW, Badhwar GD, Peterson LE, Saganti P, Dicello JF. Space radiation cancer risks and uncertainties for Mars missions. Radiat Res. 2001;156: 682–688.

5. Cucinotta FA, Kim MY, Ren L. Evaluating shielding effectiveness for reducing space radiation cancer risks. Radiat Meas. 2006; 41:173–185.

6. Cucinotta FA, Kim MY, Chappell L. Space radiation cancer risk projections and uncertainties-2012. NASA TP 2013-217375; 2013.

7. Cucinotta FA. Space radiation risks for astronauts on multiple International Space Station missions. PLoS One 2014;9(4): e96099.

8. Cucinotta FA. A New approach to reduce uncertainties in space radiation cancer risk predictions. PLoS One 2015; 10(3): e0120717.

9. Cucinotta FA, To K, Cacao E. Predictions of Space Radiation Fatality Risk for Exploration Missions. Life Sci Space Res. 2017; 13, 1–11.

10. NRC. Technical evaluation of the NASA model for cancer risk to astronauts due to space radiation. National Research Council. The National Academies Press: Washington DC; 2012.

11. NCRP. Radiation protection for space activities: supplement to previous recommendations. National Council on Radiation Protection and Measurements Commentary 23: Bethesda MD; 2014.

12. Cucinotta, F.A., Alp, M., Rowedder B., Kim, M.Y.: Safe days in space with acceptable uncertainty from space radiation exposure. Life Sci Space Res. 2015; 2: 54–69.

13. Cacao, E., Hada, M., Saganti, P.B., George, K.A., Cucinotta, F.A.: Relative biological effectiveness of HZE particles for chromosomal aberrations and other surrogate cancer risk endpoints. PLoS One 2016; 11(4): e0153998.

14. Grant EJ, Brenner A, Sugiyama H, Sakata R, Sakadane A, et al. Solid cancer incidence among the Life-span study of atomic-bomb survivors: 1958-2009. Radiat Res. 2017; 187: 513–537.

15. Leuraud, K, Richardson DB, Cardis E, Daniels RD, Gillies M, et al. Ionising radiation and risk of death from leukemia and lymphoma in radiation-monitored workers (INWORKS): an international cohort study. Lancet Haematology 2015; 2, e276.

16. Richardson DB, Cardis E, Daniels RD, Gillies M, O’Hagan JA, et al., Risk of cancer from occupational exposure to ionizing radiation: retrospective cohort study of workers in Frances, the United Kingdom, and the United States (INWORKS). Brit Med J. 2015; 351: h5359.

17. Richardson DB, Cardis E, Daniels RD, Gillies M, Haylock R, et al. Site-specific solid cancer mortality after exposure to ionizing radiation. Epidemiology 2018; 29: 31–40.

18. Preston DL, Ron E, Tokuoka S, Nishi N, Soda M, et al. Solid cancer incidence in atomic bomb survivors: 1958-1998. Radiat Res. 2007; 168: 1–64.

19. United Nations Scientific Committee on the Effects of Atomic Radiation. Sources and effects of ionizing radiation UNSCEAR 2006 report to the general assembly, with Scientific Annexes. New York: United Nations; 2008.

20. BEIR VII. Health risks from exposure to low levels of ionizing radiation. National Academy of Sciences Committee on the Biological Effects of Radiation. Washington DC: National Academy of Sciences Press; 2006.

21. National Vital Statistics Reports: United States Lifetables, 2017. Center for Disease Control, 2019; 68(7).

22. Surveillance, Epidemiology, and End Results (SEER) Program (www.seer.cancer.gov) SEER*Stat Database: Incidence - SEER Research Data, 9 Registries, Nov 2019 Sub (1975-2017) - Linked To County Attributes - Time Dependent (1990-2017) Income/Rurality, 1969-2017 Counties, National Cancer Institute, DCCPS, Surveillance Research Program, released April 2020, based on the November 2019 submission.

23. Cucinotta FA, Cacao E, Alp M. Space radiation quality factors and the delta-ray dose and dose-rate effectiveness factor. Health Phys. 2016; 110: 262–266.

24. Cucinotta FA, Cacao E. Non-targeted effects models predict significantly higher mars mission cancer risk than targeted effects model. Scientific Rep. 2017; 7: 1832.

25. Hoel DG. Comments on the DDREF estimate of the BEIR VII committee. Health Phys. 25; 108: 351–356.

26. Chang PY, Cucinotta FA, Bjornstad KA, Bakke J, Rosen CJ, D. N, Fairchild DG, Cacao E, Blakely EA. Harderian gland tumorigenesis: Low-dose and LET response. Radiat Res. 2016; 185, 449 – 460.

27. Fry RJM, Garcia AG, Allen KH, Sallese A, Tahmisian TN, Lombard LS, Ainsworth EJ. The effects of pituitarty isografts on radiation carcinogenesis in the mammary and Harderian glands of mice. In Biological Effects of Low Level Radiation Pertinent to Protection of Man and His Environment. Vol 1. 1976; 213–227.

28. Fry RJM, Powers-Risius P, Alpen EL, Ainsworth EJ. High LET radiation carcinogenesis. Radiat Res. 1985; 104: S188–S195.

29. Alpen EL, Powers-Risius P, Curtis SB, DeGuzman R. Tumorigenic potential of high-Z, high-LET charged particle radiations. Radiat Res. 1993; 88:132–143.

30. Kadhim M, Salomaa S, Wright E, Hildenbrandt G, Belyakov OV, Prise KM, et al. Non-targeted effects of ionizing radiation-implications for low dose risk. Mutat Res. 2013; 752: 84–98.

31. Illa-Bochaca I, Ouyang H, Tang J, Sebastiano C, Mao JH, Costes SV, et al. Densely ionizing radiation acts via the microenvironment to promote aggressive Trp53 null mammary carcinomas. Cancer Res. 2014; 74, 7137–7148.

32. Morgan, W. F. Non-targeted and delayed effects of exposure to ionizing radiation. Radiation-induced genomic instability and bystander effects in vitro. Radiat Res. 2003; 159, 567–580.

33. Maxwell, C. A. et al. Targeted and nontargeted effects of ionizing radiation that impact genomic instability. Cancer Res. 2008; 68, 8304–8311.

34. Lorimore SA, Coates PJ, Wright EG, Radiation-induced genomic instability and bystander effects: inter-related nontargeted effects of exposure to ionizing radiation. Oncogene 22, 7058–7069; 2003.

35. Belyakov, O. V. et al. Biological effects in unirradiated human tissue induced by radiation damage up to 1 mm away. Proc National Acad Sci USA. 2005; 102: 14203–14208.

36. Gailard S, et al. Propagation distance of the alpha-particle induced bystander effect: the role of nuclear traversal and gap junction communication. Radiat Res 2009; 171, 513–520.

37. Nagasawa H, et al. Role of homologous recombination in the alpha-particle-induced bystander effect for sister chromatid exchanges and chromosomal aberrations. Radiat Res. 2005; 164, 141–147.

38. Jain MR, Li M, Chen W, Liu T, de Toledo SM, Pandey BN, Li H, Rabin BM, Azzam EI. In Vivo space radiation-induced non-targeted responses: late effects on molecular signaling in mitochondria. Curr Mol Pharmacol. 2011; 106–114.

39. Hada M, Chappell, LJ, Wang M, George KA, Cucinotta FA. On the induction of chromosomal aberrations at fluence of less than one HZE particle per cell nucleus. Radiat Res. 2014; 182, 368–379.

40. Bettega D, Calzoolari P, NorisChiorda G, Tallone-Lombardi. Transformation of C3H10T1/2 cells with 4.3 MeV α particles at low doses: effects of single and fractionated doses. Radiat Res 1992; 131: 66–71.

41. Li Y, Wang M, Carra C, Cucinotta FA. Modularized Smad-regulated TGFβ Signaling Pathway. Mathematical Biosci. 2012; 240: 187–200.

42. Wilson JW, Townsend LW, Shinn JL, Cucinotta FA, Costen RC, Badavi FF, Lamkin SL. Galactic cosmic ray transport methods: Past, present, and future. Adv. Space Res. 1994;14: 841–852.

43. Cucinotta FA, Kim MY, Schneider I, Hassler DM. Description of light ion production cross sections and fluxes on the Mars surface using the QMSFRG model. Radiat Environ Biophys. 2007; 46:101–106.

44. Zeitlin, C. et al. Measurements of the energetic particle radiation environment in transit to Mars on the Mars Science laboratory. Science 2013; 340: 1080–1084.

45. Kim MY, Cucinotta FA, Nounu H, Zeitlin C, Hassler DM, Rafkin SCR, et al. Comparison of Martian surface ionizing radiation measurements from MSL-RAD with Badhwar-O’Neill 2011/HZETRN model calculations. J Geophys Res. 2014; On-line First. DOI: 10.1002/2013JE004549.

46. ICRP. International Commission on Radiological Protection. Assessment of radiation exposure of astronauts in space. Thousand Oaks, CA: Sage Publications; ICRP Publication 123, Ann. ICRP 42(4); 2013.

47. Dicello JF, Christian A, Cucinotta FA, Gridley DS, Kathirithhamby R, Mann J, et al. *In vivo* mammary tumorigenesis in the Sprague-Dawley rat and microdosimetric correlates. Phys Med Biol. 2004; 49: 3817–3830.

48. Weil MM, Ray FA, Genik PC, Yu Y, McCarthy M, Fallgren CM, et al. Effects of ^28^Si ions, ^56^Fe ions, and protons on the induction of murine acute myeloid leukemia and hepatocellular carcinoma. PLoS Ond. 2014;9(8): e104819.

49. Grahn D, Lombard LS, Carnes BA. The comparative tumorigenic effects of fission neutrons and Cobalt-60 γ rays in B6CF1 mouse. Radiat Res. 1992;129:19–36.

50. Imaoka T, Nishimura, Kakinuma S, Hatano Y, Ohmachi Y, Yoshinaga S, et al. High relative biological effectiveness of carbon ion irradiation on induction of rat mammary carcinoma and its lack of H-ras and Tp53 mutations. Int J Radiat Oncol Biophys. 2007;69: 194–203.

51. Trani D, Datta K, Doiron K, Kallakury B, Fornace Jr AJ. Enhanced intestinal tumor multiplicity and grade *in vivo* after HZE exposure: mouse models for space radiation risk estimates. Radiat Environ Biophys. 2010;49: 389–396.

52. Datta K, Suman S, Kallakury BV, Fornace Jr AJ. Heavy ion radiation exposure triggered higher intestinal tumor frequency and greater β-catenin activation than γ radiation in APC ^Min/+^ mice. PLoS ONE. 2013;8: e59295.

53. Illa-Bochaca I, Ouyang H, Tang J, Sebastiano C, Mao JH, Costes SV, et al. Densely ionizing radiation acts via the microenvironment to promote aggressive Trp53 null mammary carcinomas. Cancer Res. 2014;74(23): 7137–7148.

54. Wang X, Farris AB, Wang P, Zhang X, Wang H, Wang Y. Relative Effectiveness at 1 Gy after acute and fractionated exposures of heavy ions with different linear energy transfer for lung tumorigenesis. Radiat Res. 2015; 18(2): 233–239.

55. NAS, National Academy of Sciences Space Science Board. Report of the task group on the biological effects of space radiation: Radiation hazards to crews on interplanetary missions. The National Academies Press: Washington DC, 1996. 479

